# Ultrasound-Assisted Hydrolysis of Food Waste using Glucoamylase: Statistical Optimization and Mechanistic Analysis with Molecular Simulations

**DOI:** 10.1101/2023.12.14.571620

**Authors:** Avinash Anand, Karan Kumar, Kaustubh Chandrakant Khaire, Kuldeep Roy, Vijayanand Suryakant Moholkar

**Author notes:** Corresponding author: Karan Kumar, School of Energy Science and Engineering, Indian Institute of Technology Guwahati, Guwahati-781039, Assam, India.; Vijayanand Suryakant Moholkar, Department of Chemical Engineering and School of Energy Science and Engineering, Indian Institute of Technology Guwahati, Guwahati-781039, Assam, India.

## Abstract

Food waste represents a promising and cost-effective resource for synthesizing value-added products through a fermentative pathway. Preceding fermentation, the hydrolysis of food waste into monomeric sugars is a crucial step. This study presents a comprehensive investigation of food waste hydrolysis, encompassing experimental and computational approaches, using glucoamylase (GLCM) enzyme. Initial optimization of hydrolysis parameters was conducted through the Box–Behnken design of experiments, resulting in a total reducing sugar yield (TRS) of 263.4 mg/g biomass under optimized conditions within 42 hours. Sonication of hydrolysis mixture at 35 kHz at 20% duty cycle, yielded a 4× reduction in hydrolysis time with 22% enhancement in TRS yield (320 mg/g biomass). Analysis of GLCM’s secondary structure revealed sonication-induced changes through FTIR spectra deconvolution in both control and test experiments. Sonication led to a reduction in α-helix content and an increase in random coil content. Molecular dynamics simulations, including molecular docking, unveiled the majority of amino acid residues associated with the GLCM binding pocket in the α-helix and random coil regions. Consequently, sonication widened the binding pockets, facilitating easier transport of substrate and product. This effect translated into improved reaction kinetics in food waste hydrolysis.

**Research Highlights:**

- Statistical optimization of food waste hydrolysis: TRS yield = 263.4 mg/g in 42 h
- 4x reduction of hydrolysis time, 22% rise in TRS yield with 35 kHz sonication
- Sonication reduced α-helix content & increased random coil content of glucoamylase
- Molecular docking simulation to deduce mechanism of ultrasound-assisted hydrolysis
- MD simulations reveal widening of binding pockets and enhancing catalytic efficiency

## 1. Introduction

Food waste (FW) produced from diverse sources, including food processing plants, households, and commercial establishments like restaurants and cafeterias, has been investigated as a cheap and sustainable feedstock for the synthesis of value-added products such as biohydrogen, methane, bioethanol, biodiesel, enzymes, organic acids, biopolymers, and bioplastics [1–6] through fermentation route. A summary of representative literature on the synthesis of value-added products from the fermentation of food waste is provided in the supplementary material (Table S1). Typical sugar content of food waste varies from 40 to 70% (with starch content in the range 25 to 45%) depending on the type and source [7]. The starch content in food waste needs to be hydrolyzed into monomeric sugars before fermentation [5,8]. Thus, hydrolysis or saccharification is one of the important pretreatment in the fermentative synthesis of value-added products from food waste. Amyloglucosidase or glucoamylase (GLCM) has been a widely employed enzyme for the hydrolysis of starch in food waste. This enzyme exhibits a well-defined structure and function that make it indispensable for its biological activities. Structurally, GLCM is typically composed of a protein chain with a specific 3-D conformation, often organized into domains. These domains facilitate the recognition and binding of its substrate, which is typically starch or glycogen. Functionally, GLCM is primarily responsible for the hydrolysis of α-1,4-glycosidic linkages present in the polymeric chains of starch or glycogen, effectively breaking them down into smaller glucose units. This enzymatic cleavage results in the release of individual glucose molecules, which are readily absorbed and utilized by an organism as a source of energy. GLCM is widely employed in various industrial processes, including the production of high-fructose corn syrup and bioethanol, where its ability to efficiently convert starch into glucose is harnessed. In both biological and industrial contexts, structure-function relationship of GLCM underscores its significance in facilitating the breakdown and utilization of complex carbohydrates. However, in addition to the high cost of enzymes, slow kinetics is a major limitation of the enzymatic hydrolysis or saccharification of food waste [9]. Thus, intensifying the kinetics of hydrolysis by glucoamylase is crucial to the large-scale implementation of food waste-based processes for value-added products.

Sonication (or ultrasound irradiation) is known to enhance the kinetics of enzymatic processes [10–12]. Ultrasound and its secondary effect of cavitation (which essentially is nucleation, growth, and transient collapse of gas or vapor microbubbles) generate intense micro-turbulence in the liquid medium. This micro-turbulence induces changes in the secondary structures of the enzymes, which essentially leads to a rise in the activity of the enzymes. Wang et al. [13] have studied the effect of sonication on hydrolysis of starch by GLCM. The solubility of the starch improved by 136% with the application of 22 kHz sonication at an intensity of 7.2 W/mL for 10 min. Analysis of the ultrasound-exposed enzyme structure using circular dichroism revealed an increase in the random coil content of the enzyme with a concurrent reduction in α-helix and β-sheet contents. A rise in the random coil content made the structure of glucoamylase more flexible with the unfolding of proteins, which enhanced the catalytic efficiency. In another study, Meng et al. [14] studied the effect of ultrasound on the properties and conformation of glucoamylase. Moderate intensity sonication led to a rise in α-helix and random coil content of the enzyme by 17.8 and 12.4%, with a concurrent rise in the tryptophan and tyrosine population on the surface of the enzyme. These conformational changes increased the enzyme activity. High-intensity sonication, however, resulted in the deactivation of the enzyme.

The objectives of the present study are 3-fold: (1) statistical optimization of food waste hydrolysis using glucoamylase and intensification of the process at optimum conditions with sonication, (2) determination of changes in secondary structure of glucoamylase induced by sonication using deconvolution technique (FTIR), and (3) mechanistic investigation into the ultrasound-induced enhancement of glucoamylase activity with molecular docking and dynamics simulations with a representative substrate of amylotriose. Concurrent analysis of the experimental and molecular simulations results has revealed interesting mechanistic features of the ultrasonic enhancement of the food waste hydrolysis by glucoamylase (GLCM).

## 2. Materials and methods

### 2.1. Materials and initial processing of food waste

Food waste was collected from the food outlets on the institute campus. It was dried in a hot air oven at 60 °C for 72 h, followed by pulverization using a mixer grinder to a particle size of 0.85 mm (Bajaj Pluto 500 W). The pulverized food waste was stored at 25°C. Amyloglucosidase (glucoamylase) enzyme from *Aspergillus niger* was purchased from Sigma-Aldrich. All other chemicals and media components were procured from HiMedia a Pvt. Ltd., India, and used as received without any pretreatment.

### 2.2. Characterization of food waste biomass

#### 2.2.1. Determination of sugar content and composition of food waste biomass

Compositional analyses of food waste biomass were carried out by High-Performance Liquid Chromatography (HPLC) equipped with an autosampler (Shimadzu, UFLC, Prominence, Japan). The reaction mixture was prepared by adding 1 mL of 2 M trifluoroacetic acid (TFA) and 5 mg of food waste biomass to a 2 mL Eppendorf tube. The Eppendorf tube was placed in a water bath (80 °C) for two h. Hydrolyzed food waste was centrifuged at 10,000 rpm for 15 min. The supernatant was transferred to a fresh 2 mL Eppendorf tube to evaporate the remaining TFA and dried for 12 h in a hot air oven at 80 °C. TFA hydrolyzed food waste biomass was suspended in 500 µL ultrapure water followed by filtration through a syringe filter (polyvinylidene fluoride membrane) with a pore size of 0.22 µm. The monosaccharides content of TFA hydrolyzed food waste biomass was scrutinized by HPLC (Shimadzu, Model: DGU-20A5R, autosampler type) with a refractive index (RI) detector equipped with a photodiode analyzer (PDA). The HPLC employed a 300 mm × 7.8 mm Aminex@ HPX-87H column along with a guard column (catalog # 1250140, 50 mm × 7.8 mm). The reducing sugar content in hydrolyzed food waste was determined using a mobile phase of 0.05 mM H_2_SO_4_ at a flow rate of 0.6 mL/min with oven temperature maintained at 60°C. The actual sugar composition in food waste biomass was obtained using standards of five monomeric sugars, viz. maltose, glucose, galactose, arabinose, and xylose.

#### 2.2.2. Functional group analysis of food waste biomass

The functional groups in the food waste biomass were analyzed using Fourier transform infrared (FTIR) spectroscopy (Perkin Elmer, Spectrum Two, USA). Food waste biomass was mixed with potassium bromide (1:100 w/w ratio) using a mortar and pestle, and the pellets were created under a 15-ton hydraulic press. The FTIR spectra of food samples were obtained in the wave number range of 4000 – 400 cm^-1^.

#### 2.2.3. Surface morphology analysis of food waste biomass

Field Emission Scanning Electron Microscopy (FE-SEM) examined the surface morphology of food waste biomass (Carl Zeiss, Model-Gemini 300, Germany). Before placing samples of food waste biomass on the carbon tape, the carbon tape was glued to the stab surface. To prevent biomass from gaining moisture, the samples on the stab were double-gold coated in the vacuum chamber. Images of the double gold-coated samples were captured using an FE-SEM analyzer at a 5 KX magnification for surface morphology investigation.

#### 2.2.4. Thermal degradation analysis of food waste

Thermal degradation behavior (or thermogravimetric analysis) of food waste was performed with a thermal analyzer (NETZSCH, Model: STA449F3A00). The thermogravimetric (TGA) and differential thermogravimetric (DTG) curves were obtained with respect to the temperature. The sample was heated under an argon gas environment at 10 °C min^-1^ from 30 to 1000 °C.

### 2.3. Statistical optimization of enzymatic hydrolysis of food waste

The physical parameters for enzymatic hydrolysis or saccharification of food waste biomass were optimized by using the statistical 2^nd^ order Box-Behnken design (BBD) (Design Expert 9.0.7.1 version, Stat-Ease) with total reducing sugar yield per unit mass of biomass as the objective function (or response variable) [5,9]. The parameters for optimization and their ranges were: temperature (30°-50°C), citrate phosphate buffer (pH 4-6), food waste biomass loading (5-15% w/v), time of hydrolysis (12-72 h), and amyloglucosidase concentration (20-80 U/g). All experiments were repeated in triplicate to assess the reproducibility of the results. The relationship between the objective function (TRS yield) and optimization parameters was established by fitting a 2^nd^-order polynomial equation to the experimental data. Table 1 depicts the set of experiments in the Box Behnken design and their results. Experiments in the statistical design were carried out in a 5 mL test tube with a 2 mL working volume. After completion of the specific period, the reaction was arrested by immersing the test tube in a boiling water bath. The reaction mixture was centrifuged at 10000 rpm (12298 *g*) for 5 min to remove traces of particulate impurities, and the hydrolysate was separated. The total reducing sugar released in the hydrolysate was estimated using the Nelson-Somogyi method [15], while the glucose content was measured using the GOD-POD method. Finally, a validation experiment was conducted to verify the total reducing sugar and glucose yield at the predicted optimum conditions.

#### Validation experiment

The validation experiment was carried out in a 500 mL Erlenmeyer flask with a reaction volume of 100 mL. The flask was placed in an incubator shaker (Orbitek, Scigenics Biotech) operated at 200 rpm. The resulting hydrolysate was analyzed for total reducing sugar content by the Nelson-Somogyi method and the glucose content by the GOD-POD method.

### 2.4. Intensification of enzymatic hydrolysis by ultrasound

In this case, the enzymatic hydrolysis experiment was conducted in an ultrasound bath (2L Sonorex Digitec, Elma, Germany) at optimum conditions obtained using the statistical experimental design. The bath dimensions were 25 cm x 15 cm x 10 cm, and it operated at a frequency of 35 kHz with a power rating of 35 W. During the enzymatic hydrolysis reaction, sonication was performed at a duty cycle of 20%, translating to 2 minutes of sonication followed by 8 minutes of mechanical shaking for every 10 minutes of the reaction. The reaction flask was positioned at the center of the bath, submerging approximately half of its height in the water. To maintain consistent acoustic intensity throughout the bath, the flask’s position remained unchanged across all experiments. Bath water temperature was carefully controlled at 30 ± 2 °C. A control experiment involving only mechanical shaking was conducted, and 200 μL samples were periodically withdrawn from the reaction mixture to determine instantaneous total reducing sugar concentration. Control experiments continued for 42 hours, ensuring that the difference in total reducing sugar content between successive samples was less than 5%. In contrast, test experiments (including sonication) were conducted for 10 hours. Both control and test experiments were performed in triplicate.

### 2.5. Computational analysis of food waste hydrolysis by glucoamylase (GLCM)

#### 2.5.1. Selection of glucoamylase structure for molecular docking

The model system for molecular simulations comprised glucoamylase (GLCM) enzyme with alpha-1,4-maltotriose (or amylotriose), the major constituent of food waste, as the substrate. The catalytic mechanism of the GLCM enzyme was investigated using molecular docking protocols reported in our previous study [12,16]. The 3-D coordinates of GLCM from *Aspergillus niger* complexed with tris and glycerol were imported from the Protein Data Bank (PDB) [17]. Among numerous 3-D structures of GLCM available at PDB, the crystal structure with PDB ID 3EQA (1.90 Å resolution) was chosen. The docking calculations were performed using AutoDock v1.5.6 [18] integrated in MGL Tools v1.5.6. The Gaussian 09 package [19] was used to draw and optimize the chemical geometry of amylotriose. The imported GLCM had chemical moieties attached to its chemical structure, such as MAN (α-D-mannopyranose), TRS (2-amino-2-hydroxymethyl-propane-1,3 diol), GOL (Glycerol), and crystal water, which were removed before performing docking simulations. Interactions present in GLCM-amylotriose complex were analyzed in PLIP webtool (https://plip-tool.biotec.tu-dresden.de/plip-web/) and PyMOL™ v2.4.1 [20]. The docking simulations were reproduced by removing amylotriose from the GLCM-amylotriose complex, followed by redocking in GLCM to compare binding coordinates and affinities.

#### 2.5.2. Prediction of the binding pocket and active site of glucoamylase (GLCM) enzyme

Before docking experiments, the active-site of GLCM was predicted using PrankWeb (https://prankweb.cz/) in the P2Rank webserver, respectively. Later, the molecular interactions between GLCM and amylotriose and the catalytic mechanism of GLCM were investigated using molecular docking protocols as described in [16]. The chemical structure of amylotriose was drawn and optimized using Gaussian 09 package [19].

#### 2.5.3. Molecular dynamics (MD) simulations and analysis

The MD simulation analysis for validating the docking results were performed using GROMACS v2019.3 [21] software package and the analysis of MD trajectories was executed through VMD (Visual Molecular Dynamics) [22] software and its associated plugins. For detailed methodology and analysis, please refer to Appendix A1 in supplementary material.

#### 2.5.4. Analysis of the changes in the secondary structure of GLCM

We employed an FTIR-based method to explore modifications in the secondary structure of GLCM caused by sonication [23]. The secondary derivatives of peak frequencies ranging from 1600 to 1700 cm^−1^ were discerned and smoothed using a 20-point Savitzky-Golay algorithm [24]. Quantification of the multicomponent peak area under the amide-I bands of GLCM was conducted through a Gaussian function fitting program in Origin 9.0. The determination of secondary structure fractions was achieved utilizing the deconvolution methodology for FTIR spectra, as documented in previous literature [16,25].

## 3. Results and Discussion

### 3.1. Characterization of food waste biomass

The monosaccharide composition analysis of food waste biomass by TFA hydrolysis showed only two reducing sugars, glucose and maltose. FE-SEM image of the surface morphology of food waste biomass is shown in Fig. S1(A) in the supplementary material. The surface morphology of the biomass showed smooth, compact and exposed fibrous structure. Fig. S1(B) in the supplementary material shows the FTIR spectra of food waste biomass. The peaks of starch mainly occurred in the wavenumber range 3600 – 2850 cm^-1^ and 1640 – 850 cm^−1^ [26,27]. The peaks observed within the range of 3600-2850 cm^−1^ are attributed to stretching fluctuations or vibrations involving C-H or O-H bonds within the food waste biomass [28]. The stretching vibrations of C-H bonds at 2894 cm^-1^ represent the overall hydrocarbon components present in the food waste biomass. Additionally, the more prominent peaks at 3331 and 3634 cm^−1^ correspond to intra- and intermolecular O-H bond stretching, respectively. These findings provide confirmation of the presence of starch in the food waste biomass.

Both DTG and TGA analyses were used to examine the thermal decomposition and stability of food waste (refer to Fig. S1(C) in supplementary material). As per the TGA analysis, food waste biomass was stable up to 210 °C. The decomposition of food waste occurs in three stages. The first stage (75 °C to 210 °C) showed a 7% weight loss corresponding to moisture loss. The decomposition in the second step (240 °C and 450 °C) showed 58% weight loss due to fragmentation of food waste into different organic compounds [29]. In the last stage (above 450 °C) of thermal degradation, residual food waste was converted into ash.

### 3.2. Results of statistical optimization of food waste hydrolysis

As noted earlier, the optimization parameters (or variables) for enzymatic hydrolysis of food waste biomass were biomass loading (%, w/v), amyloglucosidase enzyme loading (U/g), time (h), temperature (°C) and pH, with total reducing sugar yield as the response variable or objective function. The results of the Box-Behnken experimental design are shown in Table 1. The following 2^nd^ order (quadratic) equation was fitted to the experimental data:

**Reducing sugar yield (mg/g of food waste biomass)** = -442.67 + 2.03·A + 30.95·B + 2.50·C + 10.55·D + 88.19·E – 0.003·A·B + 0.00052 A·C + 0.0016 A·D + 0.016·A·E + 0.0031·B·C + 0.0093·B·D + 0.093·B·E – 0.0016·C·D – 0.016·C·E – 0.047·D·E – 0.020·A^2^ –1.66·B^2^ – 0.028·C^2^– 0.13·D^2^ – 8.13·E^2^

Various notations are: A = glucoamylase concentration (U/g), B = biomass loading (% w/v), C = time of hydrolysis (h), D = temperature (°C), E = pH

The analysis of variance (ANOVA) of the quadratic model is given in Table 2. A large overall *F*-value (103.06) and *p*-value (< 0.0001) for the model show the significance of model terms with only a 0.01% chance of noise. The individual coefficients of the optimization variables have a *p*-value < 0.05, which indicates their significance. However, the coefficients of the interaction terms in the quadratic model have a *p*-value > 0.05, which indicates their insignificance. Physically, this result means that the optimization variables do not have any interaction among them, and their influence on enzymatic hydrolysis is essentially individual. *p*-values of model coefficients for squared variables (A^2^, B^2^, C^2^ etc.) are < 0.05, which indicates their significance. *p*-value of Lack-of-Fit is > 0.05 (insignificant), which shows that the model fits well with the experimental data. Finally, the regression coefficient of the model (*R*^2^) = 0.9861 and the predicted *R*^2^ = 0.9776 are in reasonable agreement with the adjusted *R*^2^ = 0.9766, as shown in Table 2, and also show the best fit of the model. Response surfaces for the objective function of optimization (TRS yield) between pairs of optimization variables are shown in the supplementary material (Fig. S2). The set of values of optimization variables for the highest TRS yield is pH = 5, temperature = 40 °C, biomass loading = 10% w/v, enzyme loading = 50 U/g, and time = 42 h, TRS yield (model predicted) = 255.1 mg/g biomass.

#### Validation experiment

The validation experiment of food waste hydrolysis (as noted in the previous section) was conducted at the optimized set of parameters obtained using the statistical design of experiments. The validation experiment resulted in total reducing sugar (TRS) yield of 263.4 mg/g (with glucose content of 245 mg/g). The yield of reducing sugars in the validation experiment was very close to the predictions of the quadratic model.

#### 3.2.1. Results of ultrasound-assisted hydrolysis of food waste

The total reducing sugar (TRS) yield in ultrasound-assisted food waste hydrolysis at optimum temperature conditions, enzyme concentration, biomass loading, and pH was 320.43 ± 3.8 mg/g within 10 h of total treatment. As noted earlier, the sonication was applied at 20% duty cycles, which means that the sonication period during the total treatment of 10 hours was only 2 h. We want to state that further treatment for 2 hours reduced the TRS concentration. For 11 h treatment, the TRS yield was reduced to 261.43 mg/g, and 12 h treatment further reduced the yield to 203.59 mg/g. This result is probably a consequence of the oxidative degradation of the monomeric sugar molecules induced by the radicals generated by sonication. A comparison of the results of ultrasound-assisted experiments and the control experiment (TRS yield of 263.4 ± 2.2 mg/g in 42 h of treatment) reveals that the kinetics of enzymatic hydrolysis enhanced ∼ 4×, while total reducing sugar yield increased ∼ 22 % with sonication of the reaction mixture. These results concur with the findings reported in previous literature on ultrasound-assisted enzymatic reactions [10,31].

### 3.3. Changes in the secondary structure of GLCM induced by sonication

The FTIR spectra of GLCM in control and test experiments are shown in Fig. 1A. The most sensitive region in these FTIR spectra, i.e. amide I band (1600−1690 cm^−1^) [32], which is due to C=O stretch vibration of the peptide linkage was analyzed with multi-peak fitting Gaussian function. The wavenumber ranges corresponding to various secondary structures of GLCM used for deconvolution of FTIR spectra were: β-sheets (1638 ± 2.0 cm^−1^), β-turns (1664 – 1690 cm^−1^), random coils (1648 ± 2.0 cm^−1^), α-helix (1656 ± 2.0 cm^−1^) and β-strands (1627 ± 2.0 cm^−1^) [32]. The deconvolution of the FTIR spectra of GLCM are shown in Figs. 1B and C. The percentages of different secondary structural motifs in GLCM determined from deconvolution analysis in control and test experiments are listed in Table 3.

The results presented in Table 3 reveal the significant impact of sonication on the secondary structure of GLCM. The primary change is an increase in random coil content from 28.10 % to 40.60 %, with a reduction in α-helix content from 20.95 % to 16.06 %. The content of β-sheets and β-turns also reduces with sonication.

**Table 1.** Results of the statistical experimental design for optimization of food waste hydrolysis using glucoamylase.

| Run Order | Biomass loading (% w/v) | Enzyme loading (U/g) | Time (h) | Temperature (°C) | pH | Reducing sugar yield (model, mg/g) | Reducing sugar yield (expt, mg/g) |
| --- | --- | --- | --- | --- | --- | --- | --- |
| 1 | 5 | 20 | 72 | 50 | 4 | 144.2 | 146.3 ± 3.0 |
| 2 | 15 | 20 | 72 | 30 | 6 | 151.3 | 151.6 ± 4.5 |
| 3 | 10 | 50 | 42 | 40 | 4 | 240.9 | 250.2 ± 3.1 |
| 4 | 5 | 20 | 72 | 50 | 6 | 152.7 | 145.4 ± 2.6 |
| 5 | 5 | 80 | 12 | 30 | 4 | 155.1 | 154.5 ± 4.2 |
| 6 | 10 | 50 | 42 | 40 | 5 | 255.1 | 250.0 ± 6.1 |
| 7 | 15 | 20 | 72 | 50 | 4 | 132.1 | 131.4 ± 1.0 |
| 8 | 5 | 20 | 12 | 50 | 6 | 149.2 | 152.2 ± 2.5 |
| 9 | 15 | 20 | 12 | 50 | 6 | 137.1 | 137.3 ± 3.6 |
| 10 | 5 | 20 | 12 | 30 | 4 | 145.9 | 144.4 ± 2.9 |
| 11 | 15 | 20 | 72 | 50 | 6 | 142.4 | 145.4 ± 3.2 |
| 12 | 15 | 80 | 72 | 50 | 6 | 155.3 | 155.5 ± 5.1 |
| 13 | 15 | 20 | 12 | 50 | 4 | 124.9 | 123.3 ± 3.0 |
| 14 | 5 | 20 | 72 | 30 | 6 | 163.5 | 166.5 ± 3.6 |
| 15 | 5 | 80 | 12 | 30 | 6 | 169.1 | 168.5 ± 6.5 |
| 16 | 15 | 80 | 72 | 30 | 6 | 162.3 | 161.7 ± 5.7 |
| 17 | 15 | 20 | 12 | 30 | 4 | 130.1 | 129.5 ± 5.5 |
| 18 | 10 | 50 | 42 | 40 | 5 | 255.1 | 270.0 ± 2.6 |
| 19 | 5 | 80 | 72 | 30 | 6 | 176.3 | 176.6 ± 2.9 |
| 20 | 5 | 80 | 72 | 50 | 6 | 167.4 | 170.4 ± 2.5 |
| 21 | 15 | 20 | 72 | 30 | 4 | 139.1 | 137.6 ± 2.5 |
| 22 | 10 | 50 | 12 | 40 | 5 | 226.9 | 229.1 ± 7.0 |
| 23 | 10 | 50 | 72 | 40 | 5 | 234.1 | 237.1 ± 2.0 |
| 24 | 5 | 20 | 12 | 50 | 4 | 138.9 | 138.2 ± 4.6 |
| 25 | 10 | 50 | 42 | 40 | 5 | 255.1 | 248.0 ± 4.6 |
| 26 | 15 | 80 | 12 | 30 | 6 | 153.3 | 153.6 ± 3.2 |
| 27 | 10 | 50 | 42 | 40 | 5 | 255.1 | 260.2 ± 6.0 |
| 28 | 15 | 80 | 12 | 30 | 4 | 137.3 | 139.6 ± 5.0 |
| 29 | 10 | 80 | 42 | 40 | 5 | 242.9 | 245.0 ± 3.0 |
| 30 | 5 | 20 | 72 | 30 | 4 | 153.1 | 152.4 ± 3.2 |
| 31 | 5 | 20 | 12 | 30 | 6 | 158.1 | 158.4 ± 6.1 |
| 32 | 15 | 80 | 12 | 50 | 4 | 134.0 | 133.5 ± 3.0 |
| 33 | 10 | 50 | 42 | 40 | 5 | 255.1 | 232.0 ± 7.2 |
| 34 | 10 | 50 | 42 | 30 | 5 | 245.2 | 247.3 ± 6.0 |
| 35 | 10 | 20 | 42 | 40 | 5 | 231.9 | 234.9 ± 8.9 |
| 36 | 10 | 50 | 42 | 40 | 5 | 255.1 | 268.0 ± 9.5 |
| 37 | 5 | 50 | 42 | 40 | 5 | 220.7 | 223.7 ± 10.7 |
| 38 | 15 | 80 | 72 | 30 | 4 | 148.2 | 147.7 ± 5.1 |
| 39 | 10 | 50 | 42 | 40 | 5 | 255.1 | 252.0 ± 5.6 |
| 40 | 15 | 20 | 12 | 30 | 6 | 144.1 | 143.5 ± 7.0 |
| 41 | 15 | 80 | 72 | 50 | 4 | 143.0 | 141.5 ± 6.7 |
| 42 | 5 | 80 | 72 | 50 | 4 | 157.1 | 156.4 ± 3.8 |
| 43 | 5 | 80 | 12 | 50 | 6 | 162.1 | 162.3 ± 5.5 |
| 44 | 15 | 80 | 12 | 50 | 6 | 148.1 | 147.5 ± 5.1 |
| 45 | 10 | 50 | 42 | 40 | 5 | 255.1 | 240.0 ± 4.4 |
| 46 | 5 | 80 | 12 | 50 | 4 | 149.9 | 148.3 ± 5.7 |
| 47 | 15 | 50 | 42 | 40 | 5 | 206.7 | 208.8 ± 4.1 |
| 48 | 10 | 50 | 42 | 40 | 6 | 253.1 | 249.0 ± 8.5 |
| 49 | 10 | 50 | 42 | 50 | 5 | 238.1 | 241.2 ± 4.2 |
| 50 | 5 | 80 | 72 | 30 | 4 | 164.1 | 162.6 ± 5.0 |

**Table 2.**
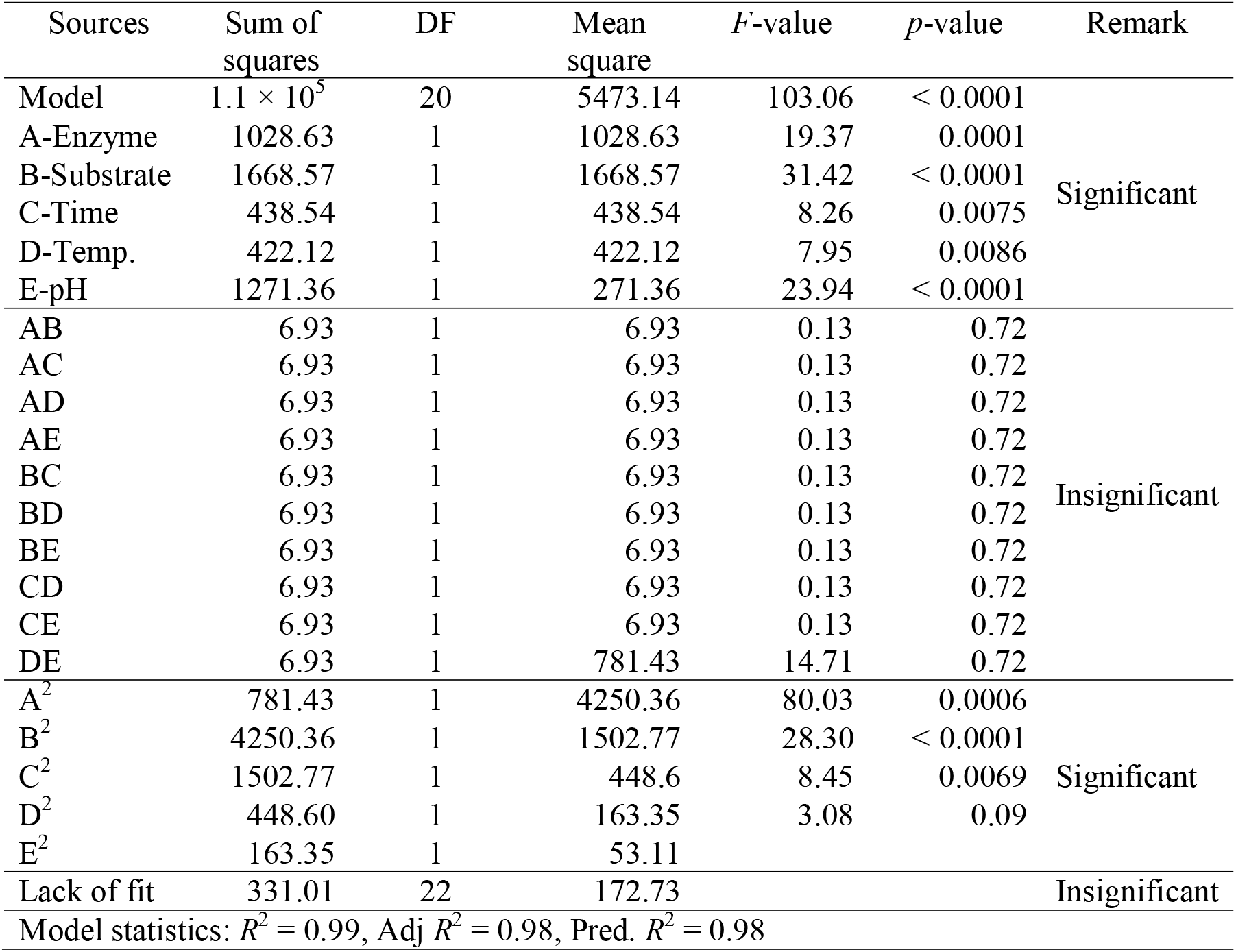
ANOVA for the statistical design of experiments.

**Figure 1.**
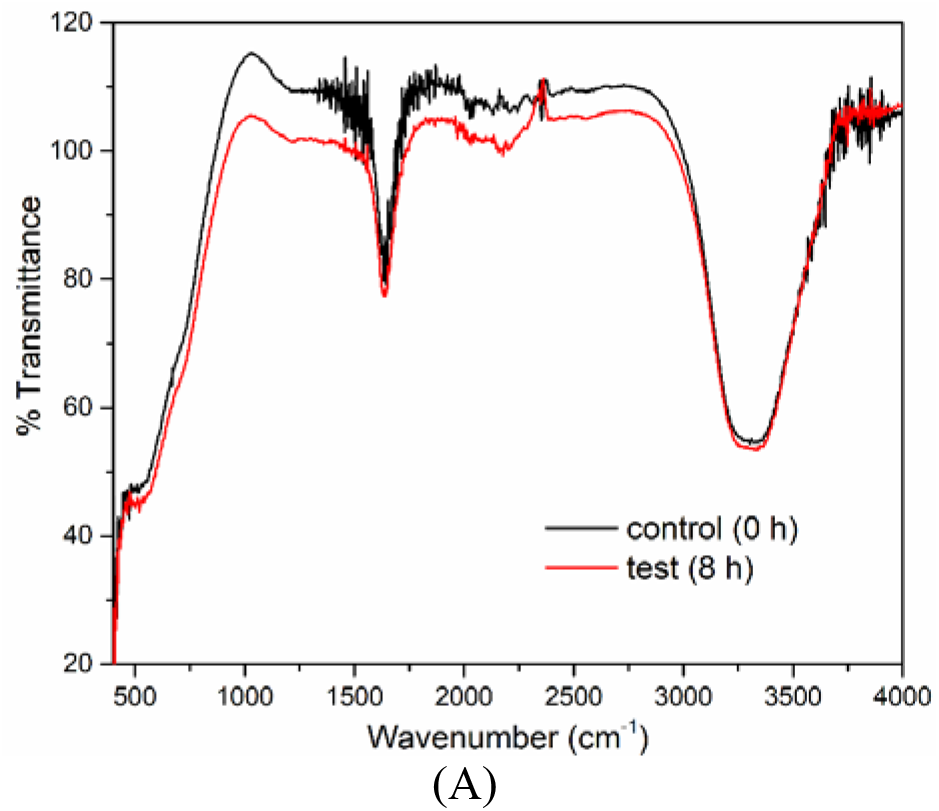
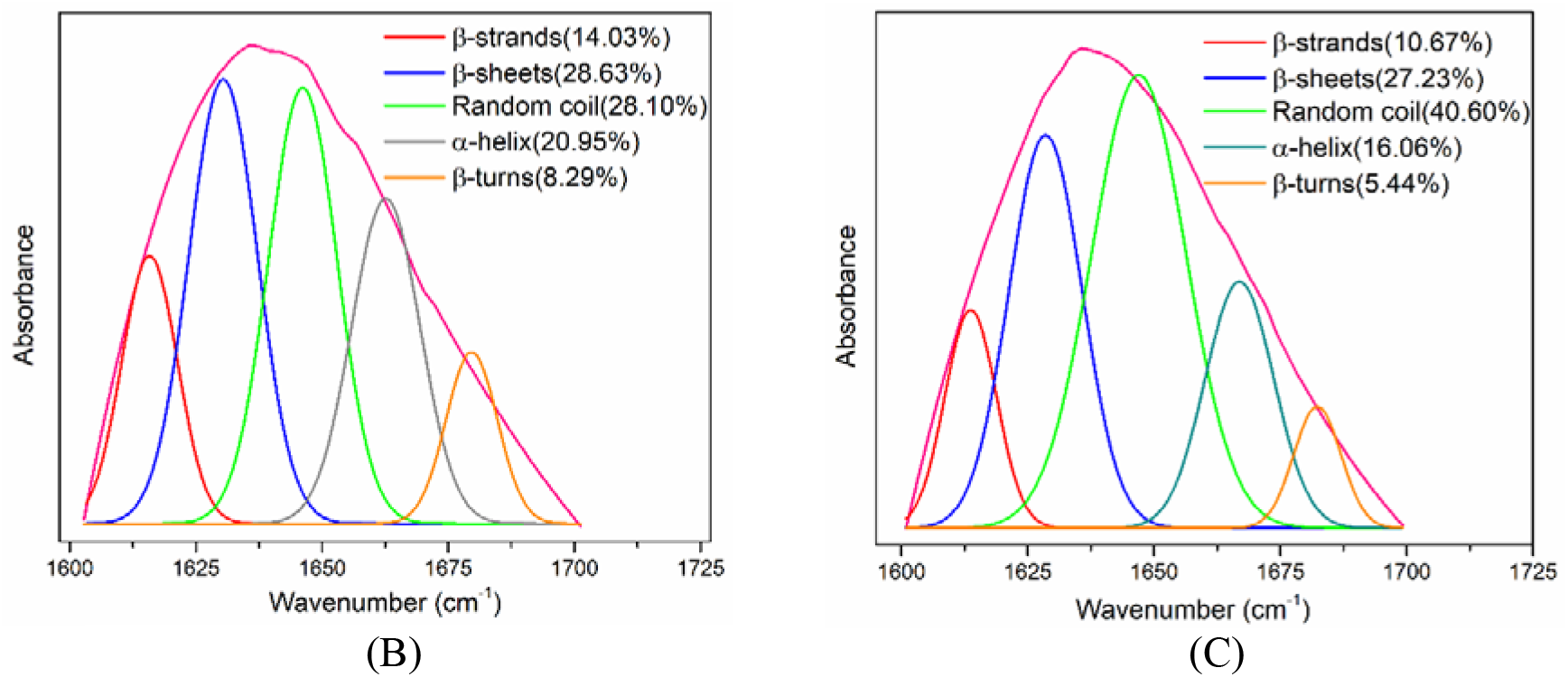
Assessing the impact of sonication on the secondary structure of glucoamylase (glcm) enzyme. (A) FTIR spectra of the glucoamylase enzyme in control conditions (hydrolysis with mechanical agitation) and test conditions (hydrolysis with mechanical agitation and sonication at a 20% duty cycle). Deconvolution of the FTIR spectra within the amide I region (1600-1725 cm^-1^) to estimate various secondary structural components of GLCM: (B) Control experiments. (C) Test experiments.

**Table 3.** Results of deconvolution of FTIR spectra for secondary structure analysis of GLCM.

| Experiment | α-Helix | β-sheets | β-strands | β-turns | Random coils | Total |
| --- | --- | --- | --- | --- | --- | --- |
| Control | 20.95 | 28.63 | 14.03 | 8.29 | 28.10 | 100 |
| Test | 16.06 | 27.23 | 10.67 | 5.44 | 40.60 | 100 |

**Table 4.** PrankWeb mediated binding pocket analysis of GLCM enzyme (PDB id: 3EQA)

| Number of binding pockets | Score | Composition of the binding pocket |
| --- | --- | --- |
| 3 | 40.85 | Chain A – ALA<sub>63</sub> PRO<sub>70</sub> ASP<sub>71</sub> TYR<sub>72</sub> TYR<sub>74</sub> TRP<sub>76</sub> ARG<sub>78</sub> ASP<sub>79</sub> LYS<sub>132</sub> ASN<sub>134</sub> TYR<sub>140</sub> THR<sub>141</sub> GLY<sub>142</sub> SER<sub>143</sub> TRP<sub>144</sub> GLY<sub>145</sub> LEU<sub>201</sub> TRP<sub>202</sub> GLU<sub>203</sub> GLU<sub>204</sub> TYR<sub>335</sub> TRP<sub>341</sub> GLU<sub>424</sub> LEU<sub>439</sub> TRP<sub>441</sub> |
|  | 7.98 | Chain A – GLY<sub>81</sub> LEU<sub>82</sub> LYS<sub>85</sub> THR<sub>86</sub> ASP<sub>89</sub> LEU<sub>154</sub> VAL<sub>215</sub> ARG<sub>218</sub> ILE<sub>277</sub> HIS<sub>278</sub> PHE<sub>280</sub> LEU<sub>343</sub> ALA<sub>347</sub> GLU<sub>350</sub> TRP<sub>441</sub> ALA<sub>445</sub> THR<sub>448</sub> ARG<sub>452</sub> PRO<sub>458</sub> |
|  | 1.23 | Chain A – LEU<sub>315</sub> LEU<sub>319</sub> ILE<sub>411</sub> THR<sub>414</sub> HIS<sub>415</sub> |

### 3.4. Molecular insight into sonication-induced enhancement of food waste hydrolysis

#### 3.4.1. Binding pocket analysis of GLCM enzyme

PrankWeb mediated binding pocket analysis of GLCM enzyme (PDB id: 3EQA) revealed the presence of 3 binding pockets in GLCM with predictability scores of 40.85, 7.98, and 1.23. Table 4 lists all three binding pockets in GLCM with their amino acid composition. This analysis revealed the predominant presence of the following amino acid residues in the binding pocket of GLCM: tyrosine, glutamine, and tryptophan. The other amino acid residues in the binding pockets are alanine, lysine, threonine, proline, and aspartic acid. The binding pockets were visualized in PyMOL, and Fig. 2 depicts the most probable binding pocket (with a score of 40.85) of GLCM. It can be seen from Fig. 2 that most of the binding pocket residues are in α-Helix and random coil of the GLCM enzyme.

**Figure 2.**
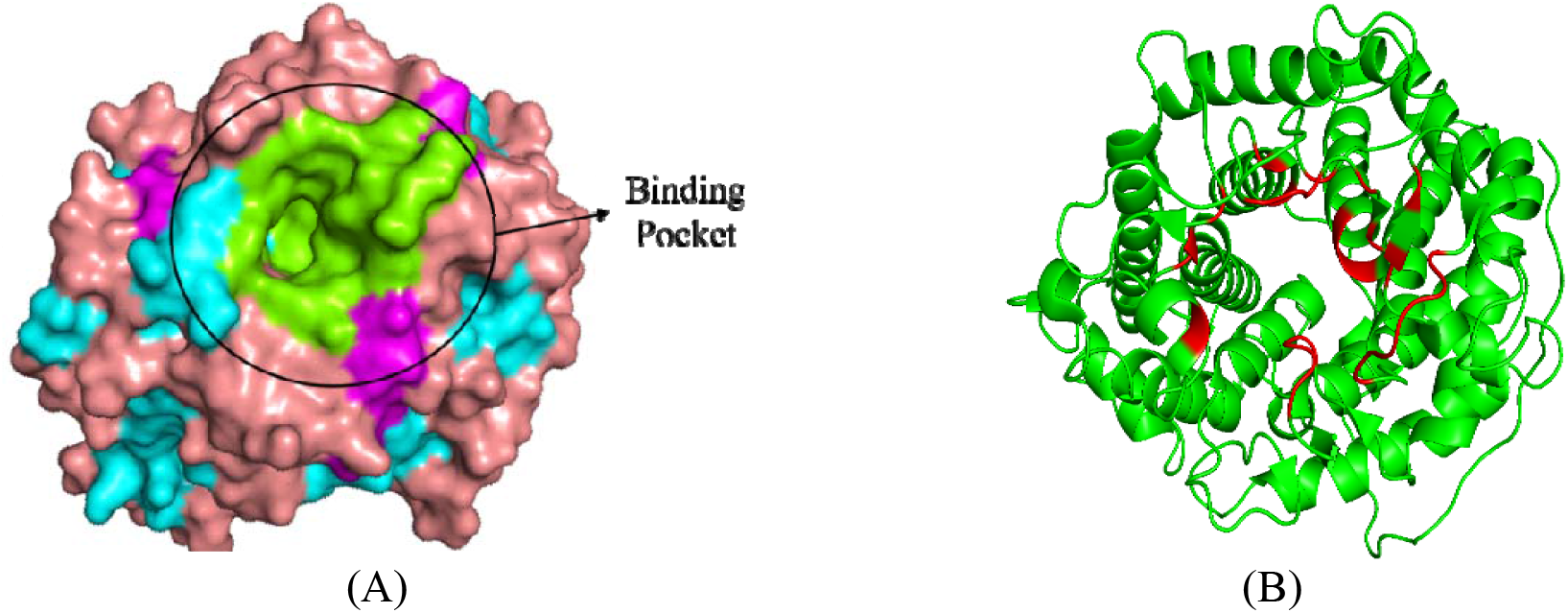
Visualization of the binding pocket of glucoamylase enzyme using PyMOL. Green-coloured residues in (A) show binding pocket residues and cavity, and red-coloured residues in (B) are residues in the most probable binding pocket (with a score of 40.85 in Table 4).

#### 3.4.2. Molecular docking simulations of amylotriose with GLCM enzyme

Molecular docking analysis was performed as described in the methodology section. The optimised structure of amylotriose (representative food waste substrate) is shown in Fig. S3 in the supplementary material. The docking analysis predicted the binding energy (ΔG) and inhibition constant (*K_i_*) of amylotriose with GLCM as -2.23 kcal mol^-1^ and 23.3 mM, respectively. The negative sign in binding energy signifies favourable binding. Moreover, docking analysis revealed that the major interactions were of hydrogen binding type. The PyMOL visualization of molecular docking of GLCM and amylotriose is shown in Fig. 3. The key residues of the GLCM involved in binding with amylotriose are ASP_71_, GLU_204_, ARG_329_, TYR_335_ and TYR_336_.

**Figure 3:**
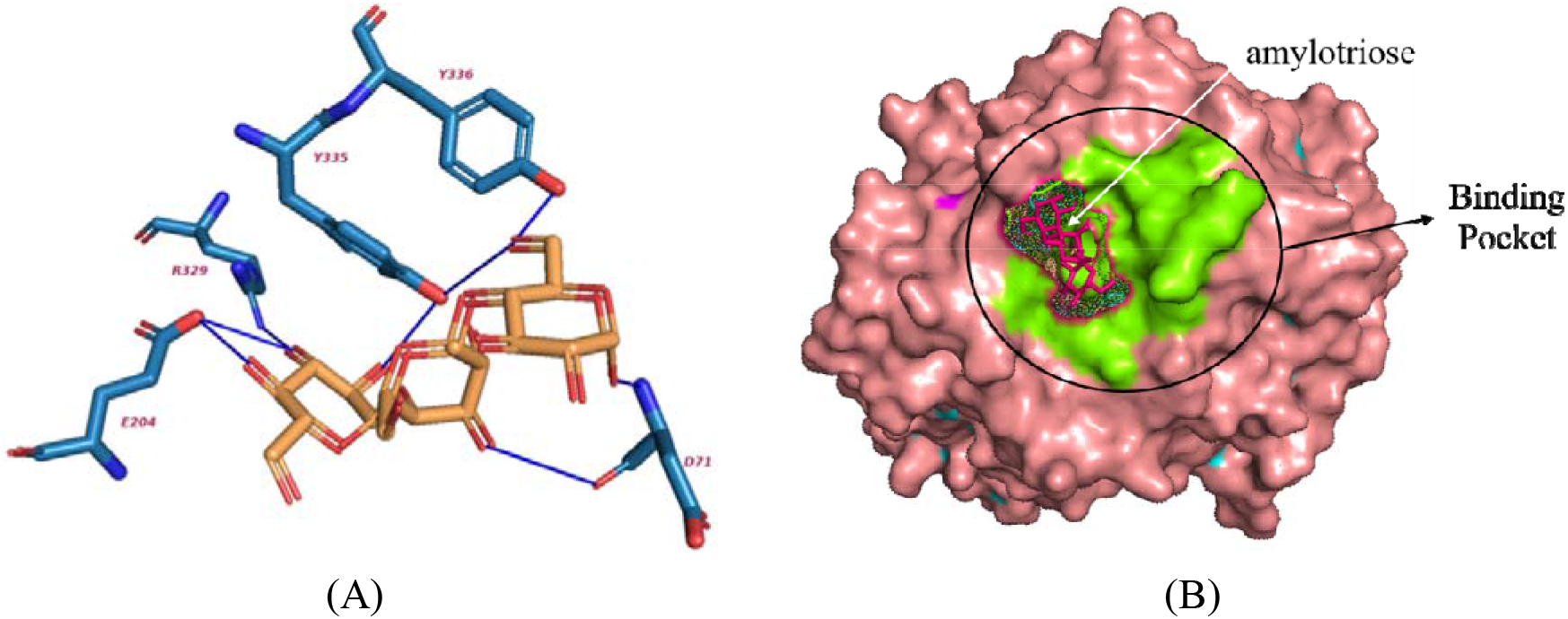
PyMOL visualization of molecular docking of GLCM and amylotriose. (A) interactions in the amylotriose-GLCM docked complex and (B) amylotriose in the binding pocket of GLCM

#### 3.4.3. MD simulation of amylotriose with GLCM enzyme

To scrutinize the influence of substrates on enzyme dynamics, a Root Mean Square Deviation (RMSD) analysis was conducted on both MD trajectories. As illustrated in Fig. 4A, the protein RMSD values from representative trajectories for each system reveal a similar flexibility of GLCM in the presence of amylotriose substrates. This observation aligns with previous findings across various systems, encompassing glycoside hydrolases and other enzymes [33–36].

##### Principal component analysis (PCA) and enzyme dynamics interpretation

To shed light on the differential effects of the amylotriose on GLCM dynamics, PCA was employed, offering insights into large protein movements. As depicted in Fig. 4B, the involvement of each residue in the primary PCA normal mode for GLCM was assessed to be essential for enzyme dynamics.

The selected mode, representing the most extensive amplitude, remained consistent in both the control simulation (APO – no substrate) and simulations with amylotriose (complex) as the substrate. However, the corresponding PCA mode varied in amplitude in complex. Fig. 4B indicates that this mode corresponds to the dynamic opening and closing of a cleft, crucial for substrate binding, release, and interaction with the catalytic center. The observations derived from the PCA mode depicted in Fig. 4C imply a more substantial interaction between GLCM and amylotriose, potentially suggesting opening of biding pocket in presence of amylotriose. Consequently, we interpret our PCA findings as indicative of a notably slower opening and closing process of the clefts surrounding the enzyme’s catalytic site when amylotripse is bound. Moreover, it is plausible that once amylotriose is bound, the anticipated opening of clefts may not occur. This dynamic movement is closely linked to catalysis, facilitating the entry of a new substrate and the release of products. Drawing from our MD simulations, we propose that higher concentration of amylotriose may, in fact, inhibit enzymatic activity (substrate inhibition). This inhibition arises from the prolonged residence of the cleaved substrate within the catalytic pocket, hindering its timely departure and consequently impeding fresh substrate molecules from accessing the reaction site. Inefficient release of reaction products, or the potential binding of a different substrate, can lead to enzyme inhibition, thereby diminishing the efficiency of biomass conversion.

**Figure 4:**
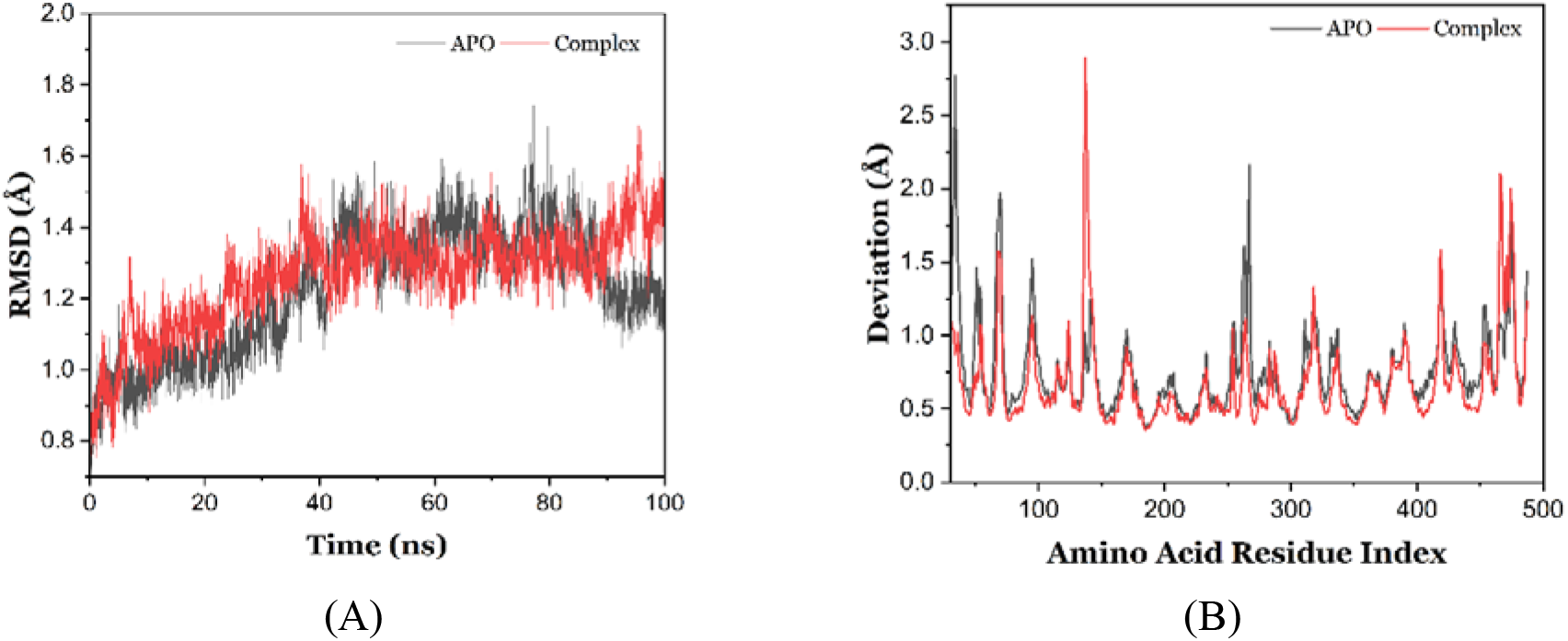
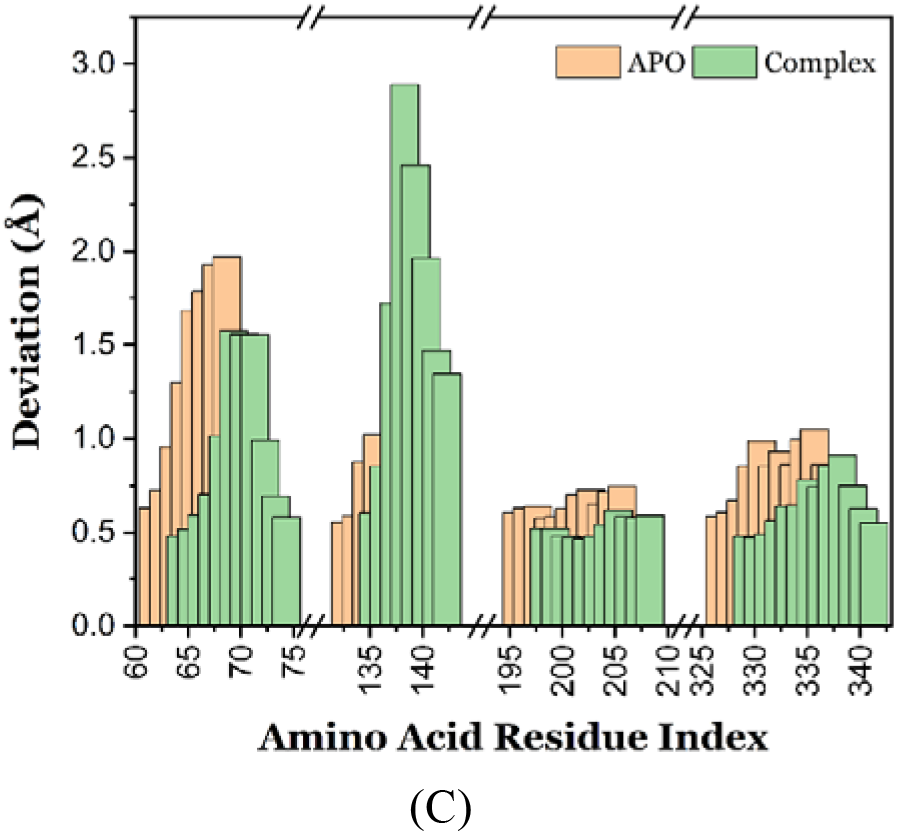
Molecular dynamics simulations and trajectory analysis of glcm in apo form and complexed with amylotriose. (A) Analysis of Root Mean Squared Deviation (RMSD) depicting structural stability in GLCM without substrate (apo, black) and with amylotriose (complex, red); (B) Individual amino acid motional amplitudes in Principal Component Analysis (PCA) for GLCM without substrate (black) and with amylotriose (red); (C) Detailed view highlighting amplitude and deviation in the primary peak region in (B), corresponding to dynamic flaps that open and close, providing access to the catalytic pocket.

#### 3.4.4. Correlation of docking analysis with modifications in the secondary structure of GLCM induced by sonication

As noted earlier, the deconvolution analysis of the FTIR spectrum of ultrasound-treated GLCM revealed a reduction in α-helix content with a concurrent rise in random coil content. As most of the amino acid residues associated with binding pockets are present in random coil and some residues are present in α-helix, the reduction in α-helix content and concurrent increment in random coil content has the following implications: (i) enzyme structure is relaxed due to unfolding of the proteins resulting in widening of the binding pockets. (ii) the relaxed enzyme structure provides easy accessibility of substrate to the binding pocket. Points (i) & (ii) essentially signify the stabilization of [E-S] or enzyme–substrate complex and improved efficiency of the enzyme, resulting in faster reaction kinetics. In Fig. 4C, the observed variation in amino acid residues within the binding pocket and catalytic residues (as detailed in Table 4) indicates a dynamic response to the presence of amylotriose, implying the opening of binding pockets. We propose that ultrasound serves to reinforce and stabilize this opening of the binding pocket, allowing a greater number of substrates, such as amylotriose, to bind more effectively to the desired site. This enhanced interaction between the enzyme and substrate is speculated to positively impact the reaction kinetics. The ultrasound-induced structural changes in GLCM, particularly the sustained opening of the binding pocket, facilitate increased accessibility of the substrate to the enzyme’s active site. Consequently, this heightened accessibility contributes to an accelerated enzymatic reaction, resulting in faster kinetics during the hydrolysis of food waste. The interplay between ultrasound-induced structural modifications and the subsequent impact on substrate binding and enzyme activity underscores the multifaceted role of sonication in optimizing the efficiency of food waste hydrolysis processes.

## 4. Conclusions

The present study has investigated three facets of food waste hydrolysis using glucoamylase (1) statistical optimization of physical parameters, (2) enhancement of hydrolysis kinetics with sonication, and (3) mechanistic investigation in sonication-induced kinetics enhancement with molecular docking and dynamics simulations. Optimization of food waste hydrolysis resulted in the following optimized parameters for TRS yield of 263.4 mg/g: biomass loading = 10 % w/v, GLCM loading = 50 U/g, temperature = 40 °C, pH = 5, time = 42 h. Application of 35 kHz sonication at 20 % duty cycle resulted marked (4×) reduction in hydrolysis time (10 h) with a 22 % rise in TRS yield (320 mg/g biomass). Deconvolution analysis of the FTIR spectrum of ultrasound–treated GLCM showed major modifications in the secondary structure of the enzyme, viz., reduction in alpha helix content and rise in random coil content. The molecular docking simulation of the binding of amylotriose (representative food waste component) with GLCM revealed presence of a majority of amino acid residues associated with the binding pocket in α-helix and random coil content of GLCM. The sonication of the enzyme essentially widened the binding pocket of the enzyme, providing easier transport of substrate/product to/from the binding pocket, which manifested in 4× faster hydrolysis with a significant rise in TRS yield.

## CRediT authorship contribution statement

**Avinash Anand:** Conceptualization, Methodology, Investigation, Visualization, Validation, writing – original draft; **Karan Kumar:** Formal Analysis, Methodology, Software, Validation, Visualization, Writing – original draft, review and editing; **Kaustubh C. Khaire:** Methodology, Investigation; **Kuldeep Roy:** Formal analysis; **V. S. Moholkar:** Supervision, Writing – review and editing

## Acknowledgements

The authors acknowledge Central Instruments Facility (CIF), I.I.T. Guwahati, for the analytical facilities. One of the authors (Karan Kumar) acknowledges the Ministry of Education, Govt. of India, for financial assistance through Prime Minister’s Research Fellowship (PMRF).

---

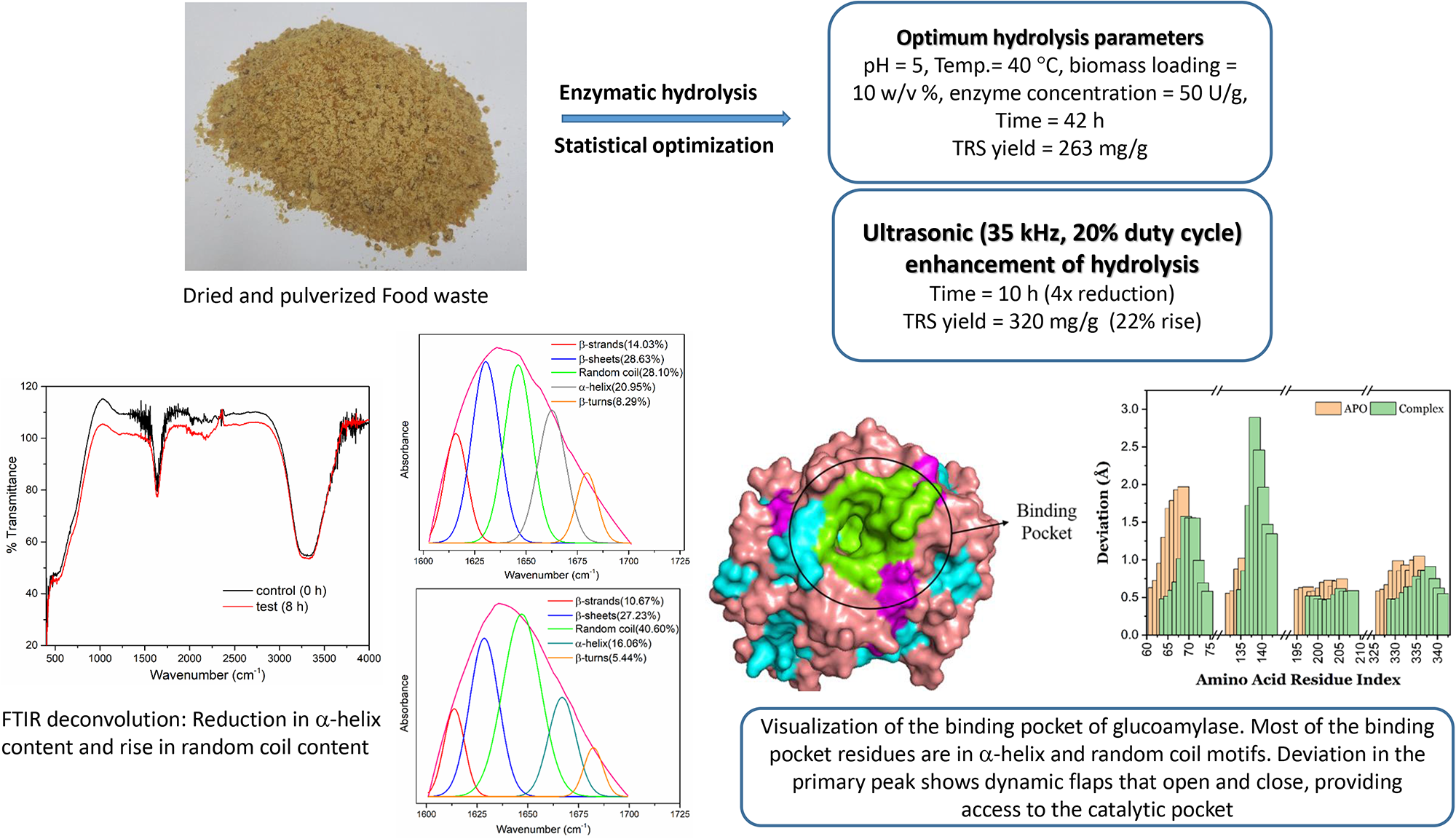

