## Supplementary material for "Ultrasound-Assisted Hydrolysis of Food Waste using Glucoamylase: Statistical Optimization and Mechanistic Analysis with Molecular Simulations"

**Appendix A1: Molecular dynamics simulations and analysis**

The study employed Molecular Dynamics (MD) simulations, employing the GROMACS v2019.3 software package for computational investigations (Abraham et al., 2015). In accordance with established methodologies, we utilized the CHARMM36-July2021 force field (Vanommeslaeghe et al., 2010) to characterize the molecular systems under scrutiny. Employing periodic boundary conditions, we maintained the systems within the NVT ensemble, ensuring a constant temperature of 310 K, and the NPT ensemble, maintaining a pressure of 1 atm. To manage temperature, Langevin dynamics were applied. Short-range non-bonded interactions were subject to a distance cutoff of 11.0 Å, while the particle-mesh Ewald (PME) method (Darden et al., 1993) was employed to calculate long-range electrostatic interactions. Integration of the equations of motion followed the r-RESPA multiple-time-step scheme, with van der Waals interactions updated every two steps and electrostatic interactions every four steps. The integration time step was uniformly set to 2 fs for all simulations within the study. In the control simulation, lacking a substrate, we initiated a 2 ns equilibration phase by gradually raising the temperature from 0 K to 310 K. Backbone atom restraints were imposed during the initial equilibration period. For simulations involving substrates, a nuanced equilibration procedure was undertaken, featuring 3 ns of backbone atom constraints followed by an additional 15 ns where backbone atoms within 5 Å of the substrate were constrained. Each system underwent rigorous MD simulations lasting 100 ns.

***Trajectory analysis***: The analysis of MD trajectories was executed through the utilization of VMD (Visual Molecular Dynamics) software and its associated plugins (Humphrey et al., 1996). To assess structural stability, we computed the Root Mean Square Deviation (RMSD) for both the ligand and the protein. Hydrogen bonds were discerned based on specific criteria: the distance between acceptor and hydrogen atoms was required to be less than 3.0 Å, and the angle formed between the hydrogen-donor-acceptor trio needed to be smaller than 30 degrees for each trajectory frame. Employing the ProDy plugin (Bakan et al., 2011), Principal Component Analysis (PCA) was conducted, affording insights into correlated atomic motions across the molecular dynamics trajectory. This analytical approach facilitated the investigation of slower motions within the enzyme's flexible regions.

**Table S1.** Value added products from fermentation of food waste

| Type of waste | Pre-treatment | Reactor type | Inoculum | Type of Biofuel | References |
|---|---|---|---|---|---|
| Food waste | Heat | 7.5L bioreactor with 3L working volume | *Seed sludge(SS)* | Biohydrogen | (Kim et al. 2006) |
| Apple pomace | Enzymatic | 150 ml bioreactor with 100 ml working volume | *HSSS* | Biohydrogen | (Wang et al. 2010) |
| FW | Heat | CSTR,500 ml working volume | *HSSS* | Biohydrogen | (Lee et al. 2010) |
| FW | Alkaline | ASBR with 0.15 m3 working volume | *HSSS* | Biohydrogen | (Kim et al. 2010) |
| FW | US with acid | Bottle with 200 ml working volume | *Seed sludge* | Biohydrogen | (Elbeshbishy et al. 2011) |

| | | | | | |
|---|---|---|---|---|---|
| FW | Lactate fermentation | Bioreactor with 150 ml working volume | *Irradiated R. sphaeriods* | Biohydrogen | (Kim et al. 2010) |
| FW | Drying | SSF, 500 ml flask | *S.Cerevisiae* | Bioethanol | (Sharma et al. 2007) |
| Waste bread | Drying | Separate,300 ml flask,80 g waste bread | *S.Cerevisiae* | Bioethanol | (Kawa-Rygielska et al. 2012) |
| FW | LAB spraying | Separate, tower shaped reactor | *S.Cerevisiae KF-7* | Bioethanol | (Tang et al. 2008) |
| FW | LAB spraying | Continuous fermenter with 4.3 kg FW | *S.Cerevisiae KF-7* | Bioethanol | (Koike et al. 2009) |
| FW | Freeze drying of waste | Anaerobic seed sludge | Two stage,UASB 8L working volume, | Methane | (Heo et al. 2004) |
| FW | Freeze drying | Anaerobic seed sludge | UASB,2.7 L working volume | Methane | (Trzcinski and Stuckey 2011) |
| FW | Fungal hydrolysis by A.oryzae | Schizochytrium mangroveri | Smf 2 L bioreactor | Biodiesel | (Pleissner et al. 2013) |
| FW | Fungal hydrolysis by A. oryzae & A. awamori, autolysis | Chlorella pyrenoidosa | Smf 2 L bioreactor | Biodiesel | (Pleissner et al. 2013) |

FW: food waste; ASBR: anaerobic sequencing batch reactor; SS:seed sludge;HSS:heat shocked seed sludge;US: ultrasonication; LAB:lactic acid bacteria;SSF :simultaneous saccharification and fermentation; Smf: submerged fermentation.

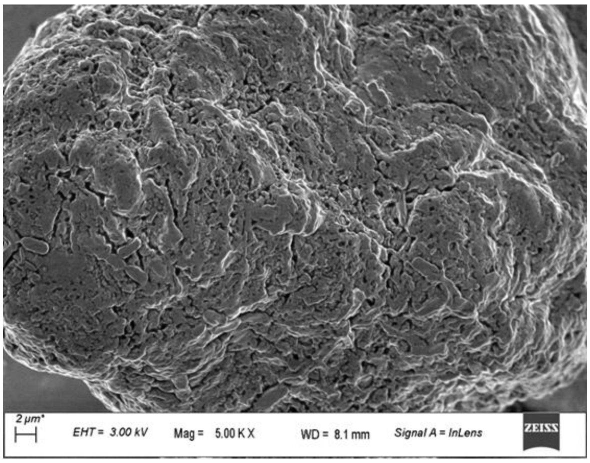


(A)

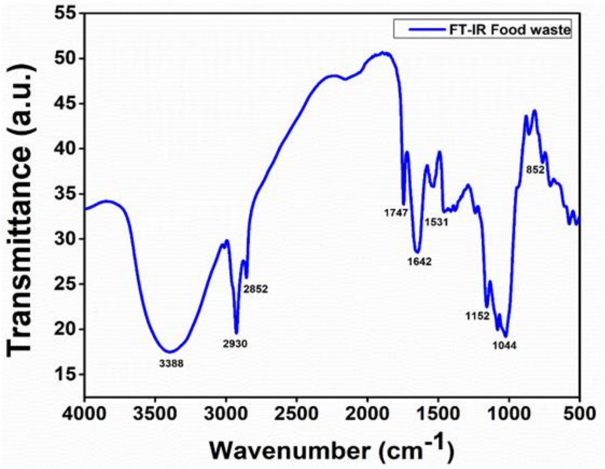


(B)

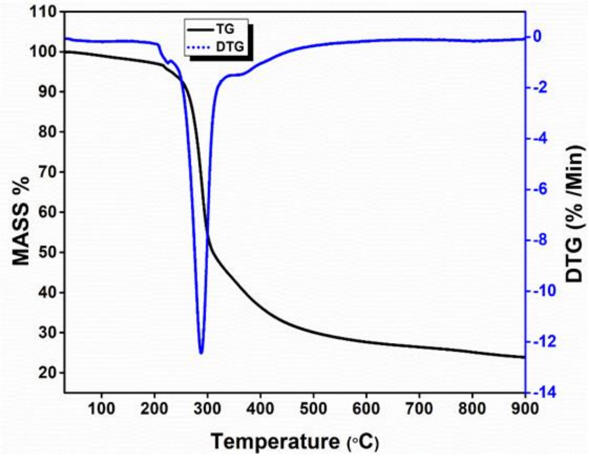


(C)

**Figure S1.** Characterization of food waste biomass. (A) FE-SEM micrograph; (B) FTIR spectrum; (C) TGA and DTG curves

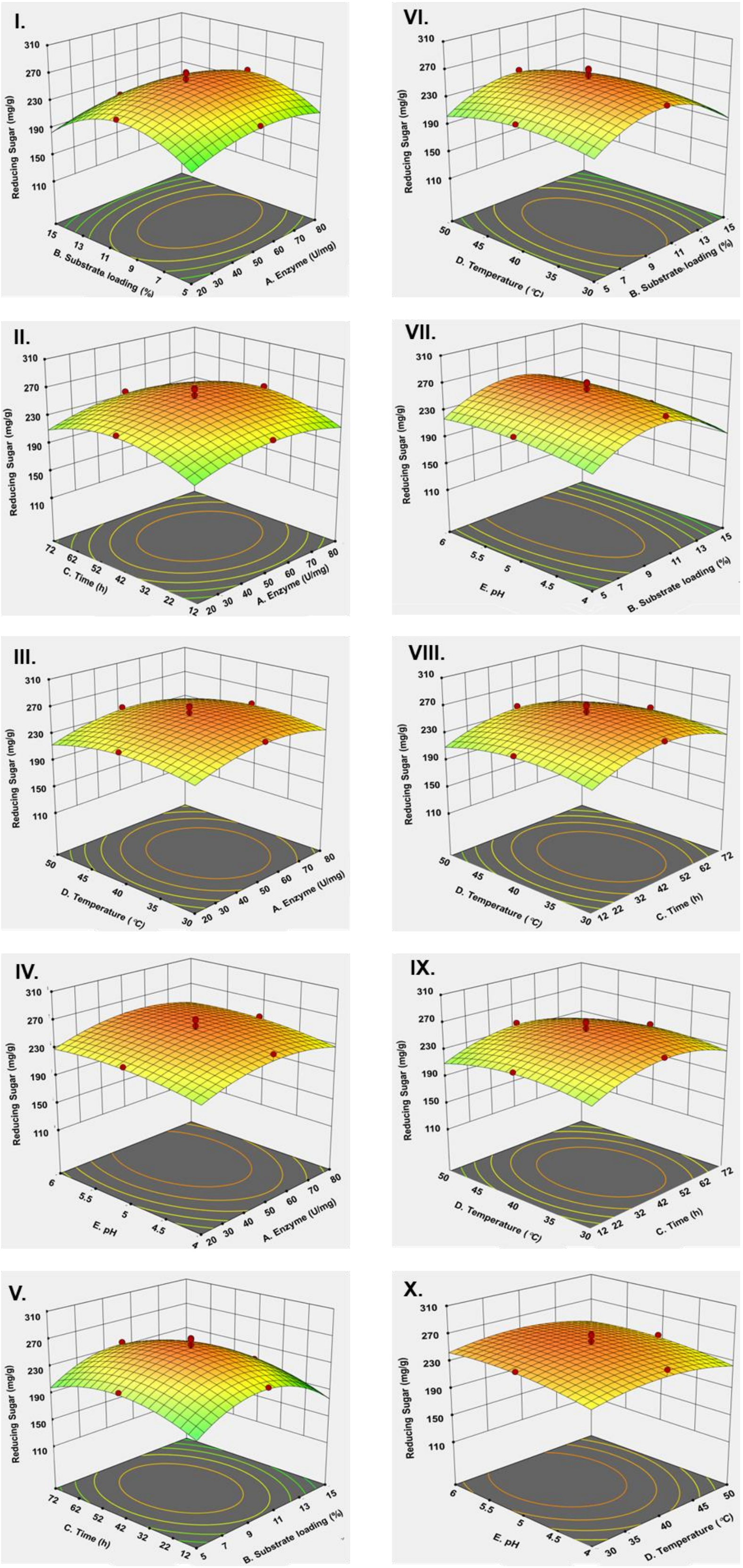


**Figure S2:** 3-D surface graphs (the interactions between independent variables) of RSM based saccharification optimization of food waste biomass in terms of reducing sugar released (mg/g). I) Enzyme loading Vs Substrate loading, II) Enzyme loading Vs Time, III) Enzyme loading Vs Temperature, IV) Enzyme loading Vs pH, V) Substrate loading Vs Time, VI) Substrate loading Vs Temperature, VII) Substrate loading Vs pH, VIII) Time Vs Temperature, IX) Time Vs pH and X) Temperature Vs pH.

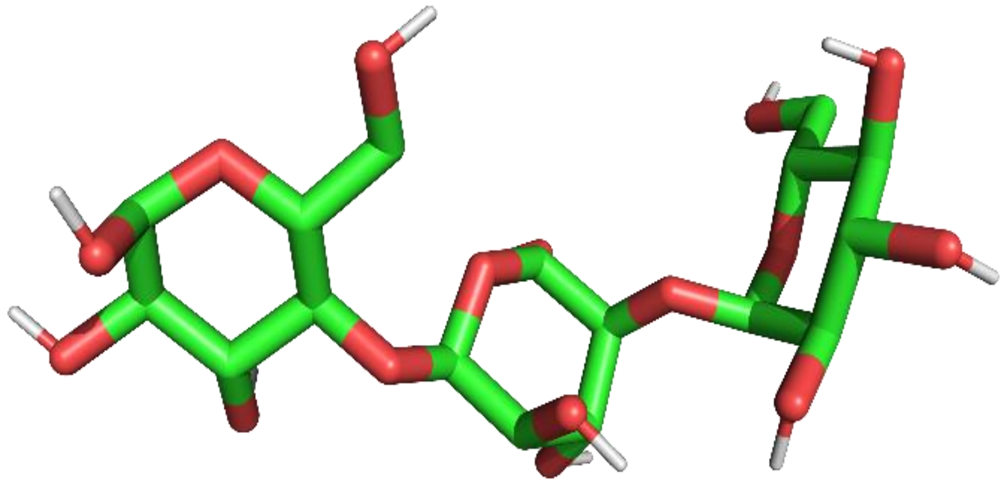

**Figure S3.** optimized structure of amylotriose.
